# Osmolarity-independent electrical cues guide rapid response to injury in zebrafish epidermis

**DOI:** 10.1101/2020.08.05.237792

**Authors:** Andrew S. Kennard, Julie A. Theriot

**Affiliations:** Biophysics Program, Stanford University, Stanford, CA 94305, USA; Department of Biology and Howard Hughes Medical Institute, University of Washington, Seattle, WA 98195, USA

## Abstract

The ability of epithelial tissues to heal after injury is essential for animal life, yet the mechanisms by which epithelial cells sense tissue damage are incompletely understood. In aquatic organisms such as zebrafish, osmotic shock following injury is believed to be an early and potent activator of a wound response. We find that, in addition to sensing osmolarity, basal skin cells in zebrafish larvae are also sensitive to changes in the particular ionic composition of their surroundings after wounding, specifically the concentration of sodium chloride in the immediate vicinity of the wound. This sodium chloride-specific wound detection mechanism is independent of cell swelling, and instead is suggestive of a mechanism by which cells sense changes in the transepithelial electrical potential generated by the transport of sodium and chloride ions across the skin. Consistent with this hypothesis, we show that electric fields directly applied within the skin are sufficient to initiate actin polarization and migration of basal cells in their native epithelial context *in vivo*, even overriding endogenous wound signaling. This suggests that, in order to mount a robust wound response, skin cells respond to both osmotic and electrical perturbations arising from tissue injury.

## Introduction

Epithelial tissues separate organisms from the outside world and bear an ever-present risk of injury, and so epithelial wound healing is a critical homeostatic process. The initial wound response in many tissues consists of rearrangements of the actomyosin cytoskeleton to form a contractile purse string that closes the wound in concert with protrusive actin structures that promote cell migration towards the injury and help cover the damaged area (Abreu-Blanco et al., 2012; Eming et al., 2014; Rothenberg and Fernandez-Gonzalez, 2019). In order to mount a wound response, cells must detect both the presence of a wound and the direction in which the wound is located; this information must be ultimately derived from changes in the cell’s environment caused by injury. Much progress has been made in identifying the *internal* signaling events which transduce environmental change into a wound response, such as waves of calcium influx, hydrogen peroxide release, and purinergic signaling (Niethammer et al., 2009; Razzell et al., 2013; Xu and Chisholm, 2011; Yin et al., 2007; Yoo et al., 2012). Yet the precise *external* changes that induce these signaling cascades have been challenging to disentangle.

Non-keratinized epithelia exposed to aqueous environments—such as the mucosal surfaces of the human body or the skin of aquatic organisms—devote considerable energy to regulating ion transport between the environment and internal body fluids. This ion transport maintains the distinct composition and osmolarity of interstitial fluid relative to the surrounding environment: for example, interstitial fluid in saltwater fishes is hyposmotic relative to the environment, while in freshwater fishes it is hyperosmotic (Boisen et al., 2003; Potts, 1984). Differential transport of various ions can lead to charge separation across the epithelium, which generates the so-called transepithelial potential (TEP) between the two sides of the epithelial layer; this TEP is related to but distinct from the transmembrane potential, and it is sensitive to the composition of the solutions on either side of the epithelium (Dietz et al., 1967; Ussing and Zerahn, 1951). If the tissue is damaged, fluid intermixing disrupts the normal ion gradients and leads to a variety of osmotic and electrical changes in the environment of the epithelium, including osmotic shock and short-circuiting of the transepithelial potential.

Both the osmotic and electrical changes could act as early cues of injury. In *Xenopus laevis* (clawed frog) and *Danio rerio* (zebrafish) larvae, the wound response is inhibited when the composition of the external medium resembles that of interstitial fluid (Fuchigami et al., 2011; Gault et al., 2014), but this observation alone cannot distinguish between osmotic and electrical mechanisms. In zebrafish epidermal cells, cell swelling due to osmotic shock following injury has been shown to provide a physical, cell-autonomous cue of tissue damage, and this cue is amplified and relayed to other cells with subsequent extracellular ATP release (Gault et al., 2014). Electrical currents have been measured emanating from wounds in many animal tissues, including rat cornea, tails of *Xenopus* tadpoles, and bronchial epithelia of rhesus macaques, and disruption of these currents has been associated with impaired healing (Reid et al., 2009, 2005; Sun et al., 2011). At the cellular level, osmotic and electrical cues both promote cell migration: hypotonic shock can promote formation of lamellipodia (Chen et al., 2019) and can intrinsically stabilize a polarized actin cytoskeleton by increasing mechanical feedback through membrane tension (Houk et al., 2012). At the same time, almost all motile cells migrate directionally in the presence of an electric field, either toward the anode or toward the cathode depending on the cell type (Allen et al., 2013). These observations suggest that both osmotic and electrical changes induced by injury of epithelial tissues could promote a migratory wound response by disrupting epithelial ion transport. Crucially, the osmotic and electrical mechanisms for sensing tissue damage are physically intertwined, and it is unclear how each signal distinctly contributes to the wound response in aqueous environments.

Due to their optical transparency and ease of experimental manipulation, zebrafish larvae have been an important model system for understanding the rapid sequence of events following tissue damage, in particular the response of epidermal cells to injury in the tailfin (Franco et al., 2019; Gault et al., 2014; Mateus et al., 2012; Yoo et al., 2012). The zebrafish larval epidermis is bilayered, with a superficial cell layer originating from the enveloping layer in early embryogenesis, and a basal cell layer that resides on a collagenous basal lamina and is specified by the *ΔNp63* promoter (Bakkers et al., 2002; Le Guellec et al., 2004; Rasmussen et al., 2015; Sonawane et al., 2005). Because zebrafish are freshwater organisms, the osmotic gradient across the zebrafish epidermis is large: external culture medium has an osmolarity of about 10 mOsm/l while the osmolarity of interstitial fluid inside the fish is estimated to be about 270-300 mOsm/l (Boisen et al., 2003; Gault et al., 2014). This gradient is maintained by a variety of ionocytes that span across the two epidermal cell layers (Guh et al., 2015).

Previous work in zebrafish has shown that, within seconds after injury, the basal cell layer reacts to tissue damage primarily by active cell migration while the superficial layer reacts by purse string contraction around the wound (Gault et al., 2014). The speed of the wound response in this tissue implies that equally rapid environmental changes must initiate this process. Although osmotic changes have been identified as one wound response cue (Gault et al., 2014), osmotic changes alone lack directional information and can only signal that a wound has occurred; they must be combined with other cues to determine the *direction* of the wound in relation to any individual cell. Electrical perturbations accompanying a drop in external osmotic pressure would in principle provide a natural directional cue to guide cell polarization and migration in the wound response, but the role of electric fields in guiding migration *in vivo* in the first few minutes after injury has not been explored.

Here we focus on the cues that specifically initiate the cell migration behaviors associated with the early wound response in the zebrafish epidermis. Through live imaging of the actin cytoskeleton in basal cells immediately following injury, we observe a range of differences in the initial wound response under different environmental conditions. We report that, in addition to the ‘osmotic surveillance’ mechanism for wound detection previously identified, zebrafish epidermal cells are also sensitive to ion-specific cues following tissue damage, independent of osmolarity and cell swelling. These ion-specific cues are consistent with expected changes in electrical activity at wounds, and in support of this mechanism we show that electric fields are capable of guiding cells and overriding endogenous wound cues, suggesting that disruption of the electrical properties of tissues may be an important injury signal in the zebrafish epidermis.

## Results

### Tissue laceration induces a rapid and coordinated wound response

A variety of wounding techniques have been used to observe the injury response in the zebrafish tailfin, including tail transection with a scalpel, laser wounding, and burn wounding (Gault et al., 2014; Miskolci et al., 2019; Yoo et al., 2012). We were specifically interested in the migratory response immediately following tissue damage, and so we developed a wounding technique—which we refer to as tissue laceration— which led to a strong and reproducible early migratory response to injury. In our laceration approach, a glass rod is pulled to a fine point, and the tissue is impaled with the needle at locations dorsal and ventral to the terminus of the notochord. The needle is dragged in a posterior direction through the surrounding tissue, tearing the tail fin (Fig. 1 A).

**Figure 1.**
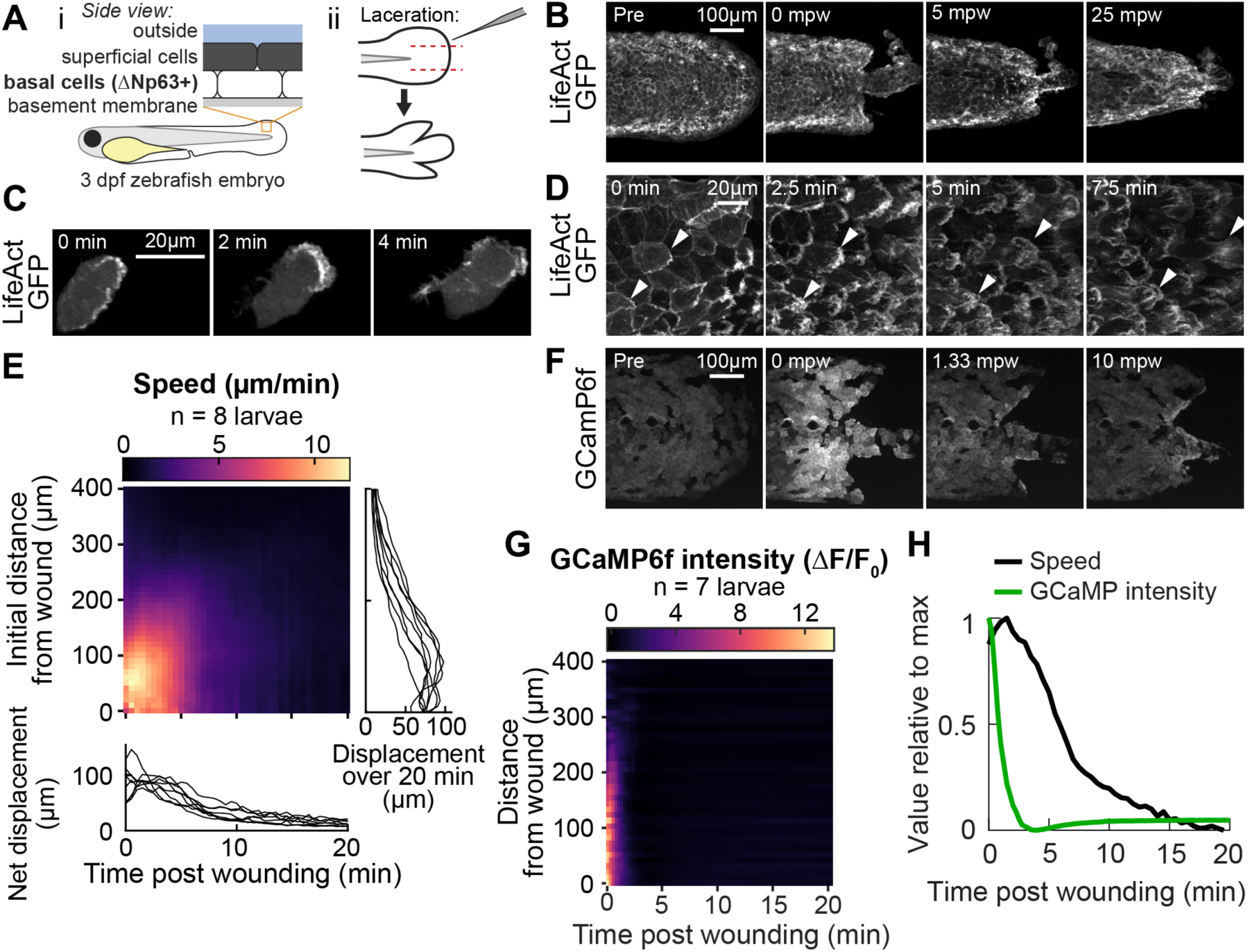
**Tissue laceration induces a rapid and coordinated wound response** (A) Schematic of (*i*) bilayered larval zebrafish skin and (*ii*) laceration technique. (B) Lacerated tailfin over time from a larva 3 days post fertilization (dpf) expressing LifeAct-EGFP in basal cells (*TgBAC(ΔNp63:Gal4); Tg(UAS:LifeAct-EGFP); Tg(hsp70:myl9-mApple))*. mpw: minutes post wounding. (B-F) are all maximum intensity Z projections of spinning-disk confocal images. (C) Individual cell from 3 dpf larva expressing LifeAct-EGFP mosaically in basal cells (*TgBAC(ΔNp63:Gal4)* larva injected with *UAS:LifeAct-EGFP* plasmid at the 1-cell stage). Wound was to the right approximately 1-2 minutes earlier. (D) Cells in a lacerated tailfin over time from 3 dpf larva expressing LifeAct-EGFP in basal cells (*TgBAC(ΔNp63:Gal4); Tg(UAS:LifeAct-EGFP); Tg(*hsp70*:myl9-mApple))*, approximately 1-2 minutes post wounding. Arrowheads: examples of individual actin-rich protrusions are followed over time. (E) Kymograph indicating speed of basal cells at a given distance from the wound over time, (N = 8 larvae). Line graphs show net displacement over space (*right*) and time (*bottom*) for each individual larva. See *Methods* and Fig. S1 for details of motion tracking analysis. (F) Lacerated tailfin from larva expressing GCaMP6f in basal cells (*TgBAC(ΔNp63:Gal4)* larvae injected with *UAS:GCaMP6f-P2A-nls-dTomato* plasmid at the 1-cell stage). mpw: minutes post wounding. (G) Kymograph of GCaMP6f intensity, normalized by the coexpressed nuclearly localized dTomato intensity, and relative to the normalized intensity pre-wounding (*F0*) (N = 7 larvae). (H) Line graph of normalized profiles of the speed and GCaMP intensity over time, averaged over 300 µm of tissue closest to the wound. To emphasize comparison of the temporal relationship, profiles are rescaled to lie between 0 and 1 (in arbitrary units).

We found that lacerated tissue rapidly reorganized and contracted around the wound site over a period of about 15-20 minutes (Fig. 1 B), consistent with wound closure observed with other methods mentioned above. A direct comparison with tail transection revealed similar spatial patterns of tissue rearrangement (Supp. Fig. S1 A). In timelapse videos of wounds from both techniques, laceration wounds induce a more pronounced migratory response within the first few minutes after wounding, suggesting that laceration wounds may be ideal for studying the early stages of the wound response (Supplemental Video 1). To determine whether this tissue reorganization was mediated at least in part by actin-based migration of cells in the basal layer, we investigated actin organization during wound closure using a basal cell-specific Gal4 driver fish crossed to a fish expressing LifeAct-EGFP from the UAS promoter.

Inspection of wound closure at high magnification revealed dynamic cytoskeletal rearrangements accompanying basal cell migration (Fig. 1 C, D and Supplemental Videos 2 and 3). Prior to wounding, the basal cell F-actin distribution was enriched uniformly around the cell at cell-cell junctions (Fig. 1 B, first image). Within two minutes of injury, actin polarized with significant accumulation of LifeAct-labeled filamentous structures on the wound-facing edges of the cell, and the formerly static cell boundary began to rapidly protrude and retract on a sub-minute timescale, reminiscent of actin ruffling (Fig. 1 C and Supplemental Video 2). For cells close to the wound, these dynamic actin-rich ruffles stabilized into lamellipodial sheets that protruded rapidly, causing the cells to elongate and translocate parallel to their long axis. This differs from the behavior of these cells when isolated in culture, where they have been studied extensively for their rapid and persistent migration and are often referred to as keratocytes (Lou et al., 2015). Isolated migratory basal epidermal cells typically adopt a wider, ‘canoe-shaped’ morphology and move perpendicular to their long axis with protrusions that maintain a persistent shape during migration (Keren et al., 2008). *In vivo*, the polarized actin ruffling response and cell elongation toward the direction of the wound was apparent for basal cells up to several hundred micrometers away, although these distant cells typically did not physically translocate (Supplemental Video 3). Following this initial rapidly dynamic wound response, the basal cells retracted their protrusive actin structures and their shapes gradually returned to resemble those of cells in an unwounded larva, though for roughly 30 minutes post wounding the cell-cell junctions continued to protrude and retract on a small scale. The migratory phase of the wound response process was rapid, with cells polarizing, migrating, and stopping within about 15-20 minutes after injury.

To better compare complex migratory behavior among many larvae and across distinct experimental conditions, we measured the local speed of cells in the basal cell monolayer over time. To do this, we developed a velocimetry analysis pipeline based on tracking the movement of many computationally detected feature points (see Supp. Fig. S1 C and *Methods*). The speed of these feature points was locally averaged in space and time to reveal the coordinated wave-like propagation of cell speed originating at the wound and traveling away (Fig. 1 E), consistent with studies of other wound types (Gault et al., 2014). Interestingly, laceration prompted a stronger cell migration response within the first 5 minutes of injury compared with tail transection (Fig. S1 B), and the profile of cell speed in space and time was reproducible across larvae despite the variation in the shapes of the laceration-induced wound margins as compared to other wounding protocols (Fig. 1 E, line graphs).

We wondered if laceration might induce transient increases in cytoplasmic calcium concentration, which have been observed with other wounding techniques (Antunes et al., 2013; Enyedi et al., 2016; Razzell et al., 2013; Xu and Chisholm, 2011; Yoo et al., 2012). To test this we injected embryos carrying the basal cell Gal4 driver with a *UAS:GCaMP6f-P2A-nls-dTomato* plasmid to express the calcium indicator GCaMP6f and a nuclear-localized dTomato fluorescent protein mosaically in basal cells. Consistent with observations from other model systems and wounding methods, we found intense and rapid propagation of increased calcium levels throughout the tailfin (Fig. 1 F). This calcium wave propagated about 5 times faster than the cell migration wave (200 µm/min vs. 40 µm/min, Fig. 1 G), implying that there is no uniform time delay between the calcium transient in a cell and the onset of migration. The same trend is observed when directly comparing the average changes in cell migratory speed and in GCaMP6f fluorescence intensity over time, normalized to comparable dimensionless quantities, where the change in GCaMP6f intensity is much faster than the change in cell speed (Fig. 1 H). This suggests that cell movement is not directly triggered or regulated by the increase in calcium, though calcium may indirectly promote wound-induced migration by functioning as a permissive cue.

Taken together, our observations of cell migration following laceration injury demonstrate a stronger migratory response in the first few minutes compared to tail transection, with overall tissue reorganization and calcium dynamics comparable to those induced by other wounding techniques. With the laceration method, we observed prominent actin-rich lamellipodia and waves of calcium and cell migration that propagated outward from the wound site at dramatically different rates.

### The wound response is sensitive to external sodium chloride, independent of osmolarity

Next we sought to determine how different physical cues might initiate the wound response in our laceration injury model. Previous work had shown that the wound response in zebrafish epidermis was inhibited by isosmotic environments (Gault et al., 2014). We confirmed this result by immersing larvae in typical freshwater medium (E3, osmolarity ∼12 mOsm/l) supplemented with sodium chloride to a final osmolarity of ∼270 mOsm/l, within the range of typical zebrafish interstitial fluid osmolarity (Gault et al., 2014; Kiener et al., 2008). Larvae wounded in isosmotic sodium chloride had a markedly reduced wound response compared to larvae in E3 (hypotonic treatment), as measured by average basal cell speed over time (Fig. 2 A, compare red with black trace).

**Figure 2.**
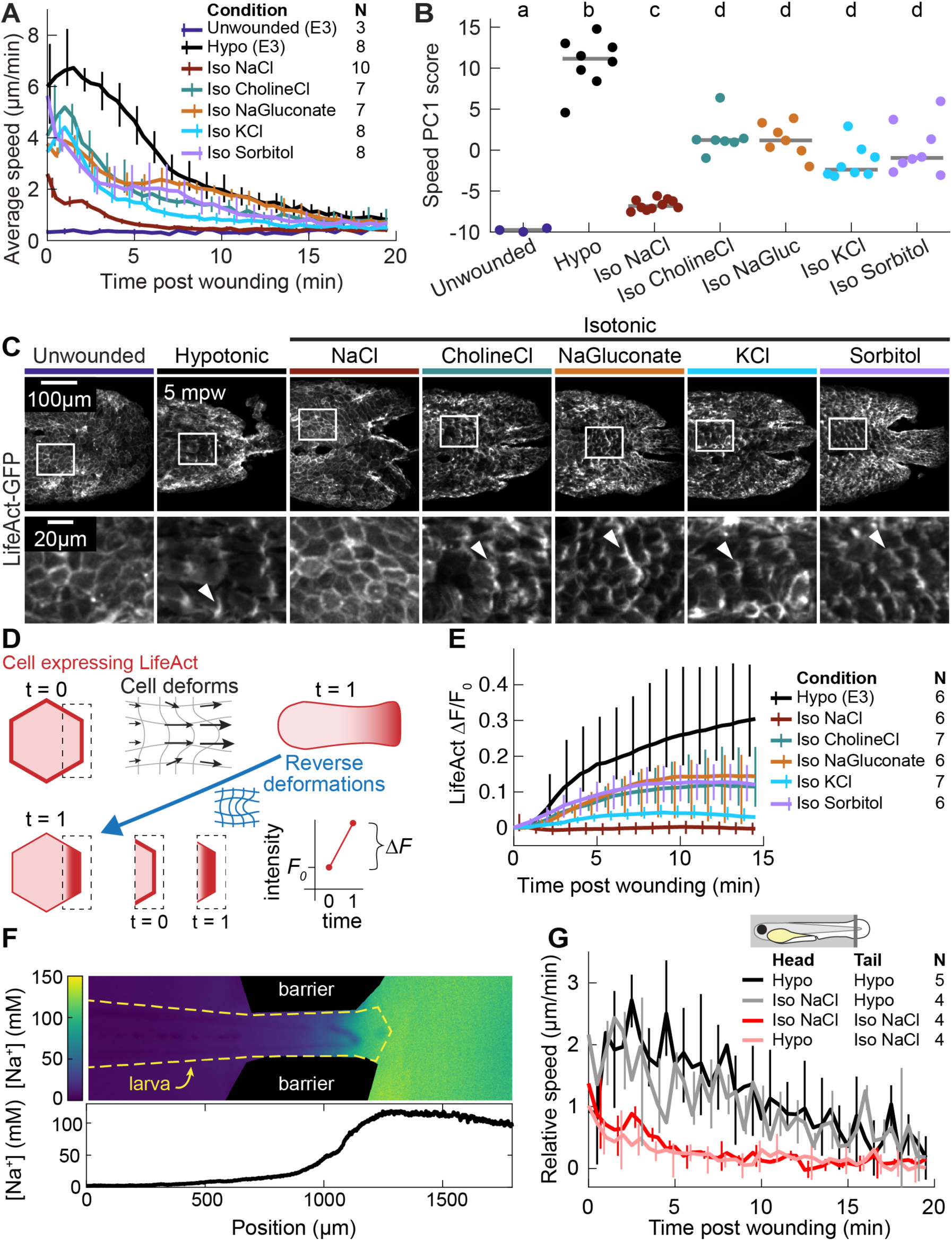
**The wound response is sensitive to local concentrations of sodium chloride, independent of osmolarity.** (A) Basal cell speed over time, averaged over 300 µm adjacent to the wound in each larva. 3 dpf larvae expressing LifeAct-EGFP in basal cells (*TgBAC(ΔNp63:Gal4); Tg(UAS:LifeAct-EGFP); Tg(hsp70:myl9-mApple))* were incubated in E3 (Hypo) or E3 supplemented with 270 mOsm/l of indicated osmolytes (Iso) and then the tailfin was lacerated and movement analyzed as described in *Methods* and Fig. S1. N indicates the number of larvae in each condition. Error bars are bootstrapped 95% confidence intervals for each condition. (B) Speed trajectories for each larva were analyzed with PCA (see Fig. S2 A – C) and each trajectory’s score along the first principal component is plotted. Grey bars indicate the mean PC1 score for that condition. Letters a-d indicate statistically distinguishable (significantly different) means (*p* < 0.001, one-way fixed-effects Welch’s ANOVA F(6, 19) = 130.9, with Games-Howell post-hoc tests). (C) (*Top*) Representative tailfins from unwounded larvae or larvae wounded in different media. Images shown from 5 minutes post wounding. (*Bottom*) Insets shown below each image. Arrowheads: examples of polarized LifeAct intensity, in the direction of the wound. (D) Schematic of computational procedure for analyzing changes in intensity, after warping image to account for cell/tissue deformation. See *Methods* for more detail. (E) Relative pixel-wise change in LifeAct intensity over time, averaged over 300 µm adjacent to the wound in each larva. (F) (*Top*) Image displaying the device allowing for different media compositions around the tailfin or the rest of the larva. Sodium concentration was calibrated with a sodium-sensitive fluorescent dye. (*Bottom*) Graph indicates the average sodium concentration along a line across the middle of the image. (G) Relative tissue speed for larvae with different media around their anterior or posterior, as shown in the diagram. To account for residual whole-larva movement due to peristaltic flow, the average tissue speed >300 µm away from the wound was subtracted from the average speed <300 µm away from the wound.

Since osmotic pressure is generated by any compounds with low membrane permeability (“osmolytes”), the osmotic surveillance model for wound detection predicts that wound response should depend only on the external concentration of osmolytes and not on their chemical identity. To test this prediction, we compared isosmotic sodium chloride treatment with isosmotic treatments of choline chloride, sodium gluconate, potassium chloride, or sorbitol. We found that, although all isosmotic treatments did reduce average cell speed, sodium chloride had a uniquely strong inhibitory effect (Fig. 2 A, compare red with other traces, and Supplementary Video 4). In contrast, the degree to which all other osmolytes inhibited a wound response was remarkably consistent with each other (Fig. 2 A, compare all other traces).

To further quantify the multifaceted differences in average migratory cell displacement over time across conditions, we treated each time profile, with 30 samples over 15 minutes, as a data point in a 30-dimensional space, and used principal component analysis (PCA) to identify the major modes of variation among these profiles. We found that over 80% of all variation could be collapsed into two principal components, which roughly corresponded to the overall amplitude of the migratory response (72% of variation) and the timing of the peak of cell movement (9% of variation), respectively (Supp. Fig. S2 A-C). When the coefficients of each profile in these first two PCA modes were plotted, profiles from unwounded larvae and larvae wounded in hyposmotic medium were situated at extreme ends of the PCA space, profiles from sodium chloride clustered near the profiles from unwounded fish, and profiles from the other isosmotic treatments fell along the continuum between the sodium chloride and the hyposmotic profiles (Supp. Fig. S2 D). The difference in magnitudes of the first principal component between sodium chloride treatment and all other treatments was statistically significant (p < 0.001, one-way fixed-effects Welch’s ANOVA F(6, 19)=130.9, with Games-Powell post-hoc tests), while the differences among the other isosmotic treatments were not statistically significant (Fig. 2 B). This quantification emphasized that the unique effect of isosmotic sodium chloride treatment on wound-induced cell migration depended on ionic chemical identity and was distinct from its osmotic effect.

Looking more closely at the dynamics of the cytoskeleton in the basal cell layer, we noticed that—with the exception of the immediate vicinity around the wound—there was a striking lack of cytoskeletal reorganization in response to wounding in isosmotic sodium chloride (Fig 2 C and Supplementary Video 4). In contrast, in all other isosmotic treatment conditions we observed transient polarization of the actin cytoskeleton in cells from the basal layer (Fig. 2 C). We quantified this observation by applying a tracking-based nonlinear warping to each image to overlay the cell intensity distributions at each timepoint, and measured the relative changes in LifeAct-GFP fluorescence intensity pixel-by-pixel across the whole tail fin over time, excluding the area approximately one cell diameter away from the wound, which was directly damaged by the laceration and would thus respond differently than cells further away (Fig. 2 D and Supp. Fig. S2 E). This analysis revealed that LifeAct intensity increased (often due to formation of lamellipodia) in all isosmotic treatments except sodium chloride, which did not induce any measurable cytoskeletal response at equivalent positions relative to the wound (Fig. 2 E). This indicates that sodium chloride’s non-osmotic effect on the wound response is associated with inhibiting cytoskeletal polarization.

It is important to note that all of these experiments were done in the presence of the fish and amphibian anesthetic Tricaine, which is a voltage-gated sodium channel inhibitor and has been shown to inhibit tail regeneration in *Xenopus* tadpoles (Ferreira et al., 2016; Tseng et al., 2010). To rule out a potential effect of Tricaine on the wound response in isosmotic sodium chloride, larvae were immobilized by injection at the one-cell stage with mRNA encoding alpha-bungarotoxin, a component of the venom of the many-banded krait snake *Bungarus multicinctus*, which immobilizes by inhibiting nicotinic acetylcholine receptors at the neuromuscular junction (Swinburne et al., 2015). Larvae treated with this alternative inhibitor were not fully immobilized, presumably due to mRNA degradation at 3 dpf, but even so the wound response in larvae immobilized by alpha-bungarotoxin in isosmotic sodium chloride was nearly identical to the response of larvae treated with Tricaine (Supp. Fig S2 F), suggesting the specific inhibitory effect of sodium chloride on the wound response is independent of the anesthetic used in the experiments.

Our findings are also robust to small variations in osmolarity: when solutions were deliberately prepared deviating from each other in osmolarity by 10% we found qualitatively similar responses in terms of migration and actin polarization (data not shown). This suggests that the unique effect of sodium chloride on the wound response is not due to small differences in osmotic strength between solutions with different ionic composition, but rather due to the actual chemical identities of the ions in solution.

### Wound response is determined by local wound environment

We next wished to determine whether the salt-specific role of sodium chloride in regulating actin reorganization during the wound response was due to local changes in sodium chloride in the wound vicinity or to global disruption of sodium chloride transport across the epidermis. Given the complex, ionocyte-mediated regulation of sodium chloride transport throughout the larval epidermis, it is possible that immersion of the entire fish in an isosmotic sodium chloride solution globally perturbs extracellular ionic composition throughout the larva. This pre-condition could lead to a general inhibition of a wound response, unrelated to location-specific cues that occur at the broken tissue barrier.

To distinguish between this global inhibition model and a model of local sodium chloride inhibition, we developed a two-chamber larval incubation device in which the tailfin was immersed in one medium and the rest of the larva in another, with the distinct media compositions maintained by peristaltic flow (see *Methods* and Fig. S2 G). Control experiments using media with different sodium chloride concentrations and a sodium-sensitive fluorescent dye as a reporter of sodium concentration confirmed that a ∼10-fold difference in sodium concentration could be maintained between the two chambers for many minutes (Fig. 2 F). When the same media was present in both chambers of the device, cell movement in response to a wound was similar to the uniform incubation conditions. When isosmotic sodium chloride media was present on only the tailfin, the wound response was identical to when the entire larva was immersed in that media (Fig. 2 G, compare red and pink traces). Moreover, when hyposmotic media was present only on the tailfin, the wound response was similar to that observed with uniformly applied hyposmotic media (Fig. 2 G, compare black and gray traces), suggesting that only the local ionic environment regulates the wound response.

### Isosmotic solutions cause comparably low cell swelling regardless of composition

What is it about the chemical composition of these media that cause them to differentially induce actin polarization and cell movement? Although we ruled out differences in *osmolarity*, it is possible that these solutions differ in *tonicity* with respect to the basal cell membrane, so that identical concentrations of different salts may differentially induce water flow across the cell membrane. To determine whether differential swelling mediates the injury signal in different isosmotic environments, we directly measured the volume of cell clusters in each condition by mosaically expressing cytoplasmic mNeonGreen in basal cells within ∼250 µm anterior of the tail fin. To measure volume, we obtained the projected area of the cluster and calculated the height at each pixel, and then integrated under this “height map” to obtain an estimate of total cell volume.

We found that basal cells near the wound swelled dramatically within 90 seconds after wounding in hyposmotic media (Fig. 3 A). To facilitate comparison across cell clusters of varying sizes, we normalized the cluster volume to the volume prior to wounding and observed the relative change in volume over time (Fig. 3 B). This revealed that basal cell clusters from larvae wounded in hyposmotic media swelled by 50% of their initial volume on average and gradually shrank over 15 minutes. In contrast, basal cell clusters from larvae wounded in isosmotic media of any composition increased in volume only slightly, with cells in sodium gluconate swelling the most—an increase of less than 6% on average.

**Figure 3.**
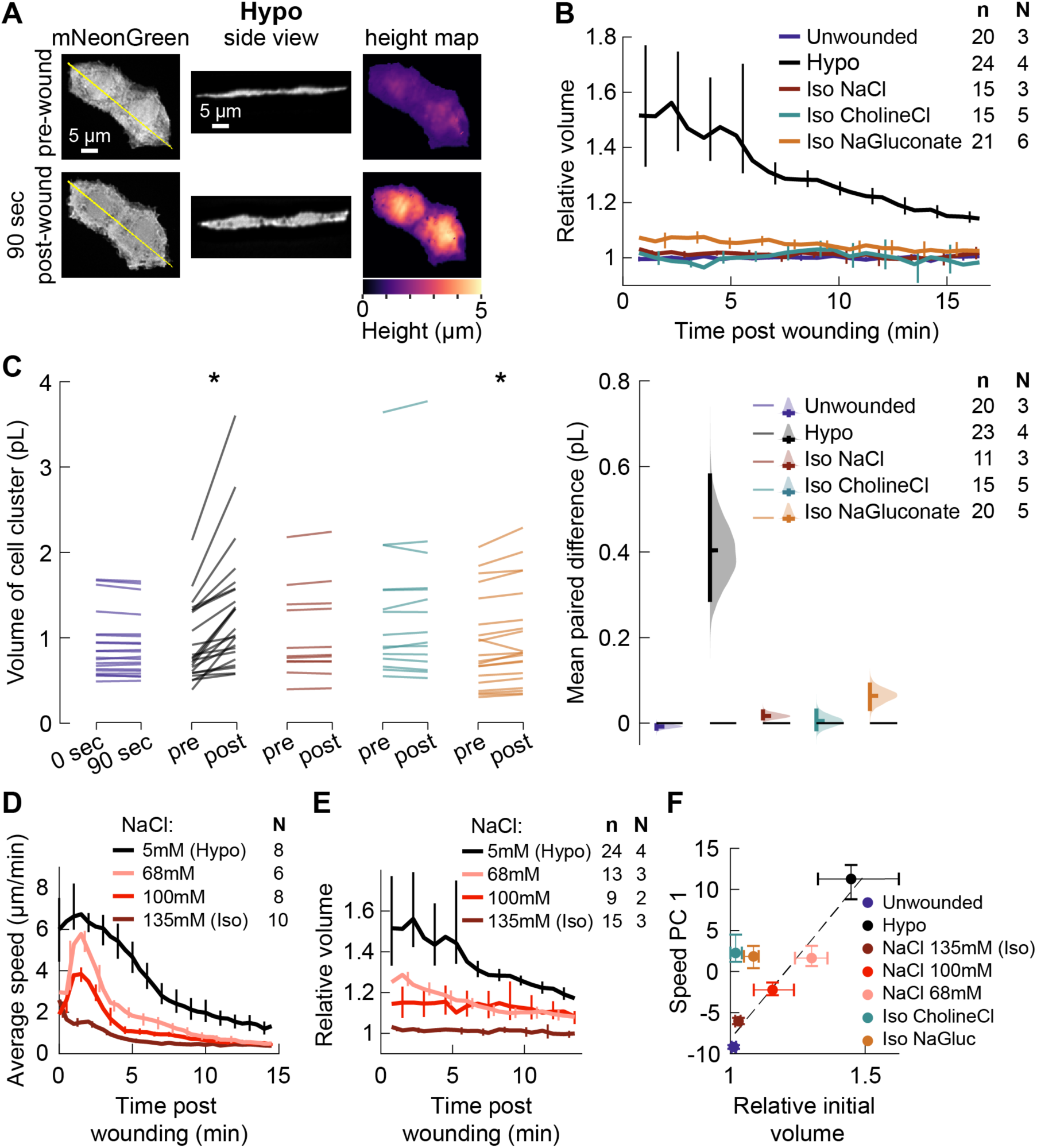
**Isosmotic solutions cause comparably little cell swelling regardless of composition.** (A) Overview of volume measurement for cell clusters. Representative cluster of cells from 3 dpf larvae mosaically expressing cytoplasmic mNeonGreen in basal cells (*TgBAC(ΔNp63:Gal4)* embryos injected with *UAS:mNeonGreen-P2A-mRuby3-CAAX* plasmid at the 1-cell stage). (*Left)* Z-projection of a representative cell cluster before and 90 seconds after wounding. (*Center*) side view at the position indicated by the yellow line. (*Right*) cell height measured at each pixel. (B) Average volume over time for cell clusters exposed to different media, relative to their volume before wounding. n: number of cell clusters. N: number of larvae. Error bars are 95% bootstrapped confidence intervals of the mean. (C) Absolute volume measurements for different cell clusters before and 90 seconds after wounding. * *p* < 0.05, two-tailed t-test on the average paired difference from each larva. (D-E) Average tissue speed (D) and cell cluster volume (E) over time for larvae treated with different concentrations of sodium chloride. 5 mM and 135 mM speed data are the same as in Fig. 2 A; 5 mM and 135 mM volume data are the same as in (B). (F) Cell cluster volume 90 seconds after wounding relative to pre-wounding volume, plotted against the 1st principal component score for speed trajectories. Error bars are 95% bootstrapped confidence intervals. Linear regression of the Hypo and three NaCl conditions displayed with a dashed line (*r^2^* = 0.95).

We used a paired data estimation plot (Ho et al., 2019) to visualize the absolute change in volume from before wounding to 90 seconds post-wounding in different media conditions (Fig. 3 C). For isosmotic media containing sodium chloride or choline chloride, the cellular volume change over this time frame was not statistically significant, nor was the magnitude of volume change in these media statistically distinguishable from that for cells on unwounded fish (p < 0.05, two-sided t-tests on the paired average volume difference from each larva). While the increase in volume in both hyposmotic medium and sodium gluconate was statistically significant, the effect size in sodium gluconate was small: the mean paired volume increase between pre- and post-wounding for clusters in sodium gluconate was 0.07 pl, while the mean paired volume increase for clusters in hyposmotic media was about 0.40 pl (95% C.I. 0.29 – 0.59 pl).

To test whether such slight swelling in isosmotic media other than sodium chloride was sufficient to explain the dramatic increase in actin polarization and migration in those media, we induced a limited degree of swelling in an orthogonal manner, by wounding larvae in intermediate concentrations of sodium chloride, and measured the degree of cell swelling and migration. As the concentration of sodium chloride decreased from isosmotic, we observed more cell migration (Fig. 3 D), but also more swelling immediately after wounding (Fig. 3 E). A linear relationship (*r^2^* = 0.95) was observed between initial volume change and degree of cell migration for the four conditions in which the concentration of sodium chloride was varied (Fig. 3 F), while the conditions in which different salts were used did not follow this same linear relationship. Instead, cells exposed to isosmotic salts other than sodium chloride moved substantially more than would be expected based solely on their volume change.

### Electric fields are sufficient to induce cell migration in the absence of wound stimuli

Having ruled out differential swelling as the cause for the specific effect of sodium chloride on the injury response, we next turned to other aspects of fish physiology that are specifically affected by sodium and chloride ions and that could lead to a differential wound response in isosmotic external concentrations of sodium chloride. One such physiological cue is the lateral electric fields generated during injury by disruption of the transepithelial potential—which itself is generated by the transport of sodium and chloride ions across the skin (Fig. 4 A) (McCaig et al., 2005; Potts, 1984).

**Figure 4.**
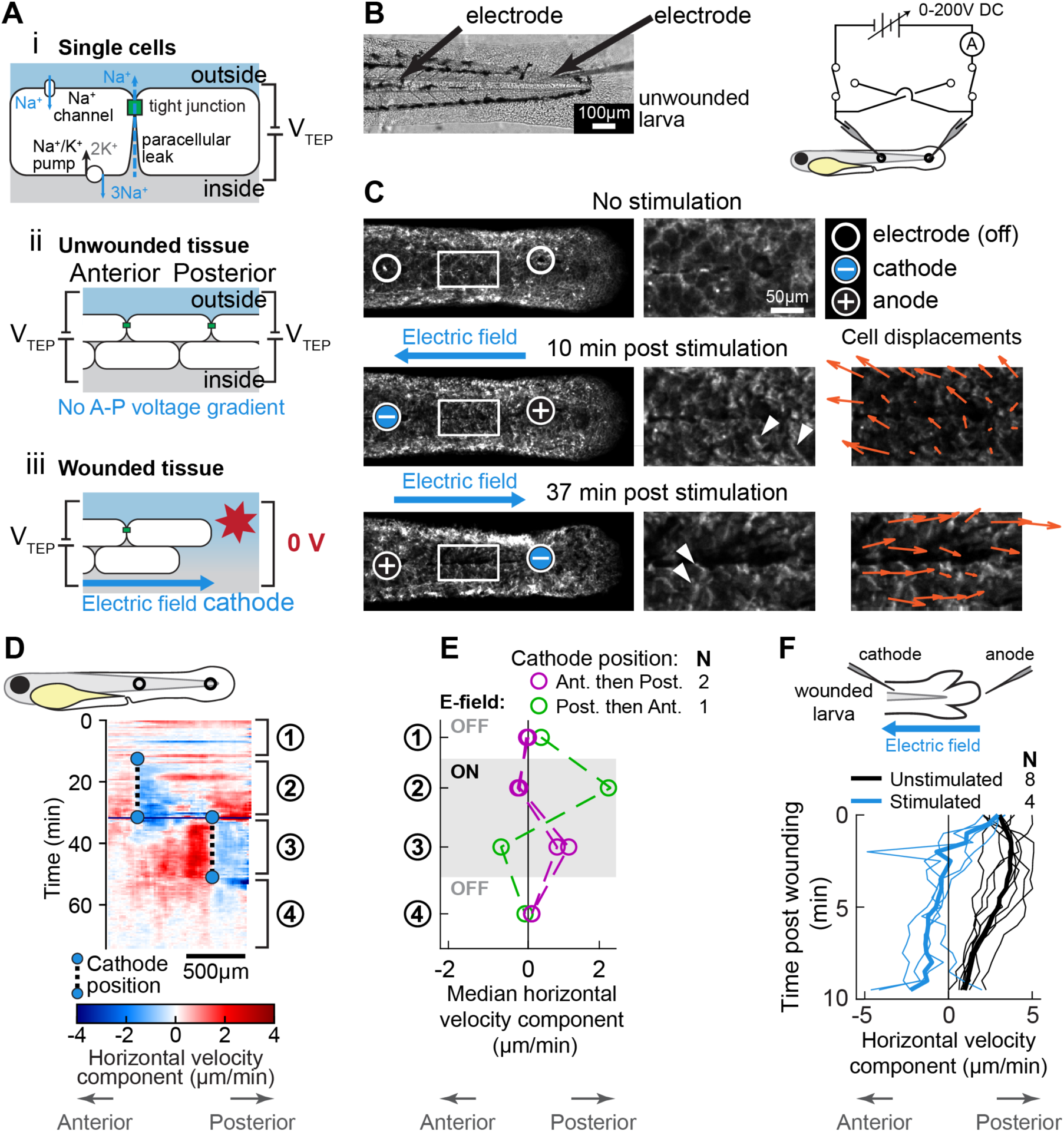
**Electric fields are sufficient to induce cell migration in the absence of wound stimuli and can override endogenous wound signals.** (A) Schematic of the origin of the transepithelial potential (TEP) due to circulating flow of sodium ions. (*ii*) Unwounded tissue does not show an anterior-posterior TEP gradient. (*iii*) Wounding short-circuits TEP leading to an anterior-posterior TEP gradient and electric field. (B) (*Left*) Brightfield image of tailfin from 3 dpf larva expressing LifeAct-EGFP in basal cells (*TgBAC(ΔNp63:Gal4); Tg(UAS:LifeAct-EGFP)),* with electrodes inserted under the skin. (*Right*) Electrical stimulation circuit with variable DC voltage, current measurement, and switches to reverse current polarity in the larva. (C) Z-projections of LifeAct signal from larva shown in (B). (*Top*) electric field off; (*Middle*) electric field on with cathode at anterior electrode; (*Bottom*) electric field reversed with cathode at posterior electrode. Stills are from one continuous timelapse. Insets are shown, and displacement vectors from tissue motion tracking are shown in orange. Arrowheads: examples of polarized LifeAct intensity oriented towards the cathode. (D) Velocity kymograph from a representative timelapse. Color indicates horizontal velocity component from tissue motion tracking analysis. Blue circles and dashed lines indicate the position of the cathode when the electric field was turned on, roughly corresponding to the empty circles on the larva diagram. Numbers 1-4 indicate different phases of the timelapse. 1: electric field off; 2: electric field on, cathode anterior; 3: electric field on, cathode posterior; 4: electric field off. (E) Median horizontal velocity component from 3 different larva. Tissue velocity was averaged in the region between the two electrodes and then the median velocity during each phase 1-4 (described above) was plotted. For one larva (shown in green) the cathode was initially positioned at the posterior electrode and was then switched to the anterior electrode. (F) Average horizontal velocity component from stimulated or unstimulated larvae. In the stimulated condition, 3 dpf larvae expressing LifeAct-EGFP in basal cells (*TgBAC(ΔNp63:Gal4); Tg(UAS:LifeAct-EGFP); Tg(hsp70:myl9-mApple))* were impaled with one electrode, with the other electrode positioned immediately adjacent to the tailfin. Larvae were then wounded and the electric field turned on, with the cathode positioned at the anterior electrode. Tissue motion between the cathode and the wound was analyzed. Thin lines represent velocity for each larva. Thick lines represent average over larvae. Unstimulated data is the same as in Fig. 2 A.

To test whether the basal layer of the epidermis will respond to electric fields *in vivo* in the absence of other wound cues, we immersed larvae in hyposmotic medium and then impaled them with two glass microelectrodes connected in series to a variable DC power supply (Fig. 4 B). After waiting 10 minutes for the minor wound response from impalement to subside, we stimulated the larvae with a DC electric field, manually maintaining a current of approximately 1 µA (see *Methods*); considering the cross-sectional area of the larval skin, this magnitude of current produces comparable current density to that previously shown to guide fish skin cell migration in culture (Allen et al., 2013). We observed that cells situated between the two electrodes rapidly polarized their actin cytoskeletons and migrated in the direction of the cathode, changing direction when the polarity of the electric field was reversed (Fig. 4 C, D, and Supplementary Video 5). The directed movement of cells toward the cathode persisted regardless of whether the cathode was initially at the anterior or posterior electrode (Fig. 4 E). suggest that, within the native tissue context, electric fields are sufficient to induce and guide cell migration.

We next tested whether cues from exogenous electric fields could dominate over endogenous wound cues *in vivo*. We impaled anesthetized larvae with a single electrode, with the second electrode placed in the media adjacent to the tailfin. Immediately after lacerating the tailfin, we turned on the electric field, with the cathode at the anterior electrode, so that the resulting electric field was the opposite polarity as compared to the field expected to be induced by injury. In all larvae tested, cells were dramatically slower in the presence of the exogenous electric field, and even moved away from the wound towards the inserted cathode, which was never observed in unstimulated larvae (Fig. 4 F). This demonstrates that, in addition to guiding cells in the absence of wound stimuli, exogenous electric fields are sufficient to override the wound response of skin cells in a living animal.

## Discussion

Our results suggest that there are at least two distinct ways in which epidermal cells detect tissue injury through changes in their external ionic environment. First, mixing of interstitial fluid with dilute external media causes cells to swell, which prompts a migratory response, as has been previously shown (Enyedi et al., 2016; Gault et al., 2014). Second, this fluid mixing specifically reduces the concentration of sodium and chloride ions around cells, which we demonstrated is sufficient to prompt actin polarization in basal epidermal cells near wounds, independent of cell swelling or any change in environmental osmolarity.

One mechanism for cells to detect wounds that would be consistent with the observation of this osmolarity-independent effect of sodium chloride is through direct detection of electric fields arising from ion transport across the epidermis. Epithelial ion transport is critical for aquatic species, which typically live in aqueous solutions of vastly different ion concentrations than their internal interstitial fluid. Sodium and chloride are the predominant ions in both the interstitial fluids and external environments of freshwater fishes, and these ions must be actively and continually absorbed from the environment to counteract leakage through tight junctions and urine (Kirschner, 2004; Potts, 1984). While transport mechanisms for these ions vary across species and environmental conditions, a consistent theme is that transport of sodium and chloride ions may be partially interdependent but are ultimately distinct: for example, the sodium-potassium ATPase can pump sodium ions against their concentration gradients, while chloride ions can be exchanged for bicarbonate ions, the concentration of which is regulated by the actions of carbonic anhydrase and V-ATPase proton pumps that expel excess protons (Guh et al., 2015; Kirschner, 2004). The activity of these various pumps and transporters can lead to charge separation across the epithelium, which manifests as the transepithelial potential (TEP).

TEPs have been measured in a variety of freshwater fish species, including trout, goldfish, and killifish, as well as in freshwater invertebrates like crayfish (Eddy, 1975; Kerstetter et al., 1970; McWilliams and Potts, 1978; Wood and Grosell, 2008; Zare and Greenaway, 1998). Sodium and chloride transport will be influenced by both concentration gradients and the TEP, and because these are the predominant ionic species, their transport will in turn affect the steady-state value of the TEP. For example, it has been proposed that differing permeability of fish skin to sodium versus chloride would lead to differing transport rates of these ions across the skin, resulting in a so-called “diffusion potential,” which could alter the TEP depending on the concentration of sodium and chloride in the external medium (Eddy, 1975; McWilliams and Potts, 1978; Potts, 1984). Such a mechanism suggests that sodium and chloride transport would have a particularly strong influence on the TEP. And as shown in Fig. 4 A, when the skin is breached, the established TEP will be short-circuited, leading to electrical potential gradients within the skin, which generate electric fields whose orientation will depend on the relative position of the wound (Reid and Zhao, 2013).

In light of this plausible connection between sodium chloride ion transport in fish skin and electrical signaling, we have shown that an electric field is sufficient to direct cell movement *in vivo* in the absence of other wound cues, and can even override endogenous wound cues at the same timescale as the normal wound response. Electric fields are an attractive physical cue for wound detection because they are intrinsically directional, providing a mechanism to rapidly coordinate cell migration towards a wound at the spatial scales of tissue. In contrast, changes in osmolarity and cell swelling do not encode directional information; these cues can initiate a wound response but must be coupled with an additional spatial cue in order to orient cell movement appropriately.

Complementary to our electrical perturbation experiments *in vivo*, electric fields have been directly measured in many tissues during regeneration on long timescales (hours and days), and the guiding effects of electric fields on cells in culture have been investigated at short timescales (tens of minutes) (Allen et al., 2013; Iglesia and Vanable, 1998; Li et al., 2012; Nawata, 2001; Zhao et al., 2006). Electric stimulation has been explored numerous times in clinical treatments for wounds and ulcers, but the lack of a detailed understanding of the mechanism by which electric fields influence cell behavior has limited progress (Gentzkow et al., 1991; Zhao et al., 2020). Our direct observation of actin polarization in response to electric fields *in vivo* at rapid timescales of 5-10 minutes bridges the gap between detailed mechanistic studies in cell culture and functional studies in tissues, and suggests that zebrafish are an ideal model system to further interrogate how cells respond to electric fields in physiological contexts.

## Author Contributions

A.S.K. performed the experiments and analyzed the data. A.S.K. and J.A.T. conceived of the experiments and wrote the manuscript.

## Acknowledgements

We thank Darren Gilmour and Jonas Hartmann for generously sharing their expertise, training, and advice on initially establishing this model system. We are also grateful to Jeff Rasmussen and Alvaro Sagasti for sharing of fish lines and expertise concerning zebrafish skin. We thank Philippe Mourrain and David Raible for graciously hosting fish in their facility and freely sharing reagents and advice, and thank Tom Daniel and Bill Moody for helpful discussions about epithelial electrophysiology. We are particularly grateful to Matthew Footer for invaluable technical advice, and also for careful critical reading of the manuscript, along with David Raible and Prathima Radhakrishnan. Finally, we are grateful to Ellen Labuz, Christopher Prinz, Mugdha Sathe, and other members of the Theriot laboratory, as well as Anna Huttenlocher, for numerous thought-provoking discussions. A.S.K. was supported by NIGMS Training Grant T32GM008294; J.A.T. acknowledges support from the Howard Hughes Medical Institute and the Washington Research Foundation.

## Methods

### Zebrafish husbandry

Zebrafish (TAB5 background WT strain) were raised and embryos harvested according to standard procedures (Westerfield, 2007). Experiments were approved by either Stanford University or University of Washington IACUCs. Animals were reared on a 14 h light, 10 h dark cycle at 28.5°C. Animals were crossed through natural spawning, and embryos were collected within 1-2 hours after spawning. Embryos were reared at 28.5°C in E3 medium without methylene blue (5 mM NaCl, 0.17 mM KCl, 0.33 mM CaCl_2_, 0.33 mM MgSO_4_) (“E3 medium,” 2008). All experiments were performed on embryos 72 - 90 hours post-fertilization.

### Transgenic zebrafish lines

The *TgBAC(ΔNp63:Gal4)^la213^*; *Tg(UAS:LifeAct-EGFP)^mu271^; Tg(hsp70:myl9-mApple)* line was generated from a natural cross of the *TgBAC(ΔNp63:Gal4)^la213^; Tg(UAS:LifeAct-EGFP)^mu271^* line—a generous gift from Alvaro Sagasti (Helker et al., 2013; Rasmussen et al., 2015)—with the *Tg(hsp70:myl9-mApple)* line (Lou et al., 2015) by screening for fluorescence, and subsequently maintained through outcrosses to TAB5 WT fish.

### Plasmid constructs and mRNA synthesis

Plasmids for microinjection were generated using Gateway cloning into Tol2kit zebrafish expression vectors (Kwan et al., 2007). The *UAS:GCaMP6f-P2A-nls-dTomato* plasmid was generated by PCR amplification of a 2kb fragment from *AAV-EF1a-DIO-GCaMP6f-P2A-nls-dTomato*, a gift from Jonathan Ting (Addgene plasmid #51083), using primers 5’-GGGGACAAGTTTGTACAAAAAAGCAGGCTTA<u>GCCACC</u>ATGGGTT-CTCATCATCATC-3’ (introduced Kozak sequence as underlined) and 5’-GGGGACCACTTTGTACAAGAAAGCTGGGTTGCCGTCGACTTACTTGTACAGC-3’. This fragment was introduced into the Tol2kit plasmid pME using BP Clonase II and standard Gateway cloning procedures (Invitrogen). This pME plasmid was recombined with the Tol2kit plasmids p5E-UAS and p3E-polyA into Tol2kit expression vector pDestTol2CG2 to generate the final plasmid. All Tol2kit plasmids were a gift from C.-B. Chien.

To construct the *UAS:mNeonGreen-P2A-mRuby3-CAAX* plasmid, mNeonGreen (Shaner et al., 2013) was amplified from an encoding plasmid with the following primers: 5’-GGGGACAAGTTTGTACAAAAAAGCAGGCTGGatggtgagcaagggcgaggag-3’ and 5’-GGGGACCACTTTGTACAAGAAAGCTGGGTCcttgtacagctcgtccatgc-3’. Upper case indicates the attB sites for Gateway recombination, and lower case indicates homology with mNeonGreen coding sequence. Similarly, mRuby3 (Bajar et al., 2016) was amplified from an encoding plasmid with the primers 5’-GGGGACAGCTTTCTTGTACAAAGTGGTAatggtgtctaagggcgaagag-3 and 5’-GGGGACAACTTTGTATAATAAAGTTGTttacttgtacagctcgtccatgcc-3’. Following Gateway recombination into pDONR221 (mNeonGreen) and pDONR P2r-P3 (mRuby3), a P2A self-cleavage site and a CAAX membrane localization tag were added using Q5 mutagenesis (NEB) with the following primers: mNeonGreen-P2A: 5’-CAGGCTGGAGACGTGGAGGAGAACCCTGGACCTgacccagctttcttgtac-3’ and 5’-CTTCAGCAGGCTGAAGTTAGTAGCTCCGCTTCCcttgtacagctcgtccatg-3’ (Upper case represents insertions of a Gly-Ser-Gly (GSG) linker and a P2A site, respectively). mRuby3-CAAX: 5’-TGAGAGTGGCCCCGGCTGCATGAGCTGCAAGTGTGTGCTCTCCtaaacaactttattatacaaagttgg-3’ and 5’-TCAGGAGGGTTCAGCTTGCCGCCGCTGCCGCCGCCGCTGCCGCCcttgtacagctcgtc catg-3’ (Upper case represents insertion of a GGSGGGSGG linker and the CAAX tag). These plasmids were then recombined along with Tol2kit plasmid p5E-UAS into Tol2kit expression vector pDestTol2CG2 to generate the final plasmid. Plasmids containing the cDNA for mNeonGreen and mRuby3 were generous gifts from Darren Gilmour and Michael Lin, respectively.

mRNA was synthesized using the SP6 mMESSAGE mMACHINE reverse transcription kit (Invitrogen). Alpha-bungarotoxin mRNA was synthesized from the plasmid *pmtb-t7-alpha-bungarotoxin*, a gift from Sean Megason (Addgene plasmid #69542) (Swinburne et al., 2015). Tol2 transposase mRNA was synthesized from the Tol2kit plasmid pCS2FA-transposase, a gift from C.-B. Chien.

### Microinjection

Embryos were injected at the 1- to 2-cell stage, into the cell (rather than the yolk). Plasmids were injected at a concentration of 20 ng/µl, with 40 ng/µl of Tol2 mRNA—the volume of these drops was not calibrated. For alpha-bungarotoxin mRNA injections, drops were calibrated to ∼2.3 nl and 60 pg of mRNA was injected into each embryo.

### Preparation of larvae for imaging

Larvae were imaged at 3 days post-fertilization (3 dpf). One day prior to imaging, any larvae with the *hsp70:myl9-mApple* transgene were transferred from E3 at 28.5°C into 20 ml scintillation vials of E3 pre-heated to 37°C for 20 minutes before being returned to 28.5°C.

Larvae were screened for transgenes of interest in the morning of 3 dpf. Larvae were anesthetized in E3 + 160 mg/l Tricaine (Sigma part number E10521) + 1.6 mM Tris, pH 7—hereafter referred to as E3 + Tricaine. Larvae were then mounted in 35mm #1.5 glass-bottom dishes (CellVis D35-20-1.5N and D35C4-20-1.5N) with 1.2% low-melt agarose (Invitrogen) in E3 + Tricaine with the dorsal-ventral axis aligned parallel to the coverslip. Excess agarose was removed from around the tail of each larva with a #11 blade scalpel, and the incubation medium was replaced with experimental immersion medium prior to wounding and imaging.

E3 + Tricaine was the base for all experimental media. Additionally, isosmotic media was supplemented with 270mOsmol/L of the indicated component.

### Tissue wounding

Solid borosilicate glass rods 1 mm in diameter (Sutter Instruments) were pulled into a needlepoint with a Brown-Flaming type micropipette puller (Sutter P-87). After the unwounded larva was imaged for several frames, timelapse acquisition was paused and the needle was maneuvered by hand to impale the larva at a position just dorsal (or ventral) to the posterior end of the notochord (see Fig. 1 A). The needle was then dragged posteriorly through the tailfin to tear the skin. This was repeated on the ventral (or dorsal) side of the notochord and then imaging was resumed. The entire procedure took 30 seconds – 1 minute.

For tail transection wounds the procedure was very similar, except a #10 blade scalpel was manually maneuvered to cut off the tail posterior to the notochord, perpendicular to the anterior-posterior axis.

### Two-chamber device experiments

Two-chamber devices were made from polydimethylsiloxane (PDMS) cast in a mold fabricated from cut acrylic, inspired by previous work (Donoughe et al., 2018; Huemer et al., 2016). Device molds were cut from extruded acrylic (McMaster) using a Dremel LC-40 laser cutter and fused with acrylic cement. A 14 mm-long piece cut from the inner portion of a 22G spinal tap needle (Beckton Dickinson, ∼375 µm in diameter) was laid across the bottom of the mold to provide a channel for positioning the larva between the two chambers, and for fluid inlet into each chamber. A diagram of the device is shown in Fig S2 G.

Prior to casting, the mold was pre-coated with 5% (w/v) Pluronic as a release agent. Sylgard 184 PDMS was mixed at a ratio of 10:1 (base : initiator), degassed, poured into the molds, degassed again, and polymerized at 50°C overnight. PDMS devices were cleaned with dish soap, sonication, and Type I water, air plasma-treated for 1 minute at 500 mTorr (Harrick Plasma PDC-001) and then immediately bonded to a #1.5 25mm round glass coverslip.

Larvae were anesthetized in E3 + Tricaine and then immobilized within the device with 1.2% low-melt agarose. This agarose was carefully removed around the tail, keeping a plug of agarose around the larva in the anterior chamber for immobilization and to prevent convective fluid mixing. Tubing for peristaltic flow was positioned using custom-built equipment (see Fig. S2 G v-vi). To maintain a stable concentration gradient, peristaltic flow was maintained in each chamber at a rate of approximately 0.3 ml/min.

To measure the concentration gradient that can be maintained in this device, solutions of E3 + Tricaine and E3 + Tricaine + 135 mM sodium chloride were prepared and supplemented with 10µM of CoroNa Green (Invitrogen Cat#C36675). A wildtype larva was mounted in the device as described above and confocal image stacks were collected over time as the two CoroNa Green containing media were flowed into either chamber of the device. From the moment at which CoroNa Green was first detectable in the field of view, it took about 12 minutes for the concentrations in both chambers to stabilize. The fluorescent intensity was converted into a sodium concentration by first background subtracting and flat-field correcting each Z-projection (Model and Blank, 2006), and then comparing pixel intensities to a fluorescence standard curve generated by imaging drops of E3 + Tricaine + 10µM CoroNa Green + sodium chloride (at different concentrations) and subjecting those images to the same intensity correction procedure. The standard curve was fit to a binding curve of the form 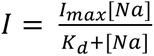 using nonlinear regression (`fitnlm` in MATLAB). The fit value of *I_max_* was 1.1529 (arbitrary units) and *K_d_* was 138 mM.

### Electrical Stimulation

Microelectrodes were pulled from thin-walled borosilicate glass capillary tubes (1 mm O.D., 0.75 mm I.D., World Precision Instruments) with a Brown-Flaming type micropipette puller (Sutter P-87) and filled with 135 mM NaCl solution. The combined series resistance of both electrodes when immersed in the same solution was ∼20 MΩ. Microelectrodes were connected into an electrical circuit using chlorided silver wires, with a variable DC power supply (PiezoDrive PD-200). Current was measured at 1 Hz sampling rate according to Ohm’s Law, by recording the voltage drop across a 100 kΩ resistor in series with the larva using a multimeter (Fluke 287).

3 dpf larvae were mounted on 25 mm #1.5 coverslips in an open bath chamber (Warner RC-40LP) and immobilized in E3 + Tricaine containing 1.2% low-melt agarose (Invitrogen). Agarose was removed from above the larvae with a #11 scalpel, leaving a thin layer of agarose surrounding and immobilizing the larva. Electrodes were maneuvered into the larval trunk with micromanipulators (Narishige MN-153). After allowing the skin to recover from impalement for 10 minutes with the circuit thus connected through the larva, the power supply was turned on.

Voltage was manually varied to maintain a current of approximately 1 µA; due to the indeterminate resistance of the electrodes + larva, the voltage required to maintain 1 µA varied between 10 - 55 V for different larvae and over the course of the experiment, as resistance slowly but steadily dropped.

For experiments combining electrical stimulation and wounding, one electrode was inserted into the larva and the other electrode was placed just outside and posterior to the larva. Tissue was lacerated as described above and the circuit was immediately turned on, with current manually maintained at approximately 1.5 µA.

### Microscopy and Image Acquisition

Images were acquired with one of two microscope setups. The first microscope used was a Leica DMI6000B inverted microscope equipped with a piezo-z stage (Ludl 96A600) a Yokogawa CSU-W1 spinning disk confocal with Borealis attachment (Andor), a laser launch (Andor ILE) with 50 mW 488 nm and 50 mW 561 nm diode lasers (Coherent OBIS), 405/488/561/640/755 penta-band dichroic (Andor), and a 488/561 dual-band emission filter (Chroma ZET488/561m). A Plan Apo 20x NA 0.75 multi-immersion objective was used. On this Leica microscope, temperature was controlled with a closed forced-air temperature-controlled heating system to maintain temperature at 28-29°C.

Alternatively, a Nikon Ti2 inverted microscope was used, equipped with a piezo-z stage (Applied Scientific Instruments PZ-2300-XY-FT), a Yokogawa CSU-W1 spinning disk confocal with Borealis attachment (Andor), a laser launch (Vortran VersaLase) with 50 mW 488 nm and 50 mW 561 nm diode lasers (Vortran Stradus), 405/488/561/640/755 penta-band dichroic (Andor), and a 488/561 dual-band emission filter (Chroma ZET488/561m). A Chroma 535/50m emission filter was also used for volume measurements and electrical stimulation measurements, where only the green channel of emission light was collected. For standard wounding experiments, an Apo 20x NA 0.95 water immersion objective was used. For volume measurements, a Plan Apo 60x NA water immersion objective was used. For electrical stimulation experiments without wounding, a Plan Fluor 10x NA 0.3 objective was used, while for electrical stimulation experiments with wounding the same 20x described above was used. On this Nikon microscope, larvae were maintained at 28-29°C using a resistive heating stage insert (Warner DH-40iL) with a temperature controller (Warner CL-100).

On both systems, full-chip 16-bit 1024×1024 pixel images were acquired with a back-thinned EMCCD camera (Andor DU888 iXon Ultra) with Frame Transfer mode and EM Gain applied. Binning was not used, except for 2×2 binning for the GCaMP data in Fig. 1 G. MicroManager v1.4.23 (Edelstein et al., 2010) was used to control all equipment, including synchronizing rapid laser line switching and piezo-z positioning with camera exposures using TTL triggers.

Two-channel z-stacks were acquired at 30-second intervals, switching laser line at each z-position before changing z-position.

### Timelapse registration

To correct for whole-body movement and drift of the tailfin, registration was performed on movies from *TgBAC(ΔNp63:Gal4)^la213^*; *Tg(UAS:LifeAct-EGFP)^mu271^; Tg(hsp70:myl9-mApple)* embryos. The myosin light chain-mApple was ubiquitously expressed, and we observed that only the skin cells migrated in response to wounding. We therefore considered myosin light chain fluorescence originating from tissues *other* than the skin to be stationary, and corrected any drift using this signal as follows. Prior to maximum intensity projection, the LifeAct intensity was thresholded and used as a mask to set corresponding regions of the myosin z-stack to 0 using custom MATLAB code; following maximum intensity z-projection, regions in the myosin channel that did not overlap with basal cells were emphasized. These modified z-projections of the myosin channel were manually cropped to select a 512×512 pixel region for registration >300 µm away from the wound. These subimages were registered in time with custom Python code by detecting KAZE features (Alcantarilla et al., 2012), matching these features between adjacent timepoints, and fitting a Euclidean transform (rotation + translation) to the feature displacement vectors using RANSAC (Fischler and Bolles, 1981). The calculated transformations were then converted to the coordinates of the LifeAct image and used to register those z-projections. Registration was performed using custom Python scripts including the following libraries: numpy (van der Walt et al., 2011), sciki-image (Walt et al., 2014), Tifffile (Christoph Gohlke, University of California, Irvine), and the python bindings for OpenCV (Bradski, 2000).

### Motion tracking and analysis

Registered LifeAct z-projections were manually aligned so the anterior-posterior axis was horizontal. Motion was tracked by detecting Shi-Tomasi corner points in each image (typically several thousand points per image) and tracking them from frame to frame using the Kanade-Lucas-Tomasi algorithm (Lucas and Kanade, 1981; Shi and Tomasi, 1994). These points correspond to areas of strong texture or curvature in the image, which make them straightforward to track. Due to high contrast and detail in the image, a majority of points could be tracked for the entire duration of a timelapse. Velocities could be calculated from the trajectories of these points.

The observation that most movement in the tailfin was in one primary direction (towards the wound) facilitated summarization of the data obtained from thousands of point tracks. To capture the most relevant tissue movement towards or away from the wound, the wounded region was manually traced, and the line between the centroid of all detected points and the centroid of the wound was computed. The positions of points (either the position in the first frame or the position in each frame) was projected onto this line to obtain a one-dimensional distance from the wound. Points were binned by their 1D coordinate along this line in 10 µm increments, and the average speed (in two dimensions) in each bin was calculated for each time point, providing a measure of velocity in one spatial dimension and time.

For the two-chamber device experiments (Fig. 2 G) overall movement of the tailfin due to peristaltic flow was not completely removed by the registration algorithm described above. To better compare measurements on different larvae, the relative velocity was used: the average velocity of points >300 µm away from the wound centroid (along the line described above) was subtracted from the velocity of each point <300 µm away from the wound centroid.

For the electric field stimulation experiments without wounding (Fig. 4 D, E), the same motion tracking approach was used, but instead of averaging the speed (the magnitude of the 2D velocity of each point), the horizontal velocity component was averaged, so that positive and negative velocities indicated movement in the anterior or posterior direction, respectively.

### GCaMP intensity tracking

Maximum intensity Z-projections were background-subtracted and manually rotated so the anterior-posterior axis was horizontal. The wound margin was manually traced, and the GCaMP6f and nls-dTomato intensities were each averaged in 10 µm increments based on the horizontal component of the displacement from the wound centroid. To correct for variation in expression, the GCaMP6f intensity in each 10 µm increment was normalized to the nls-dTomato intensity in that increment, and then *F_t_* (*x*)—the normalized GCaMP6f intensity at a horizontal position *x* and frame *t* —was further normalized to report relative changes in intensity over time, using the formula 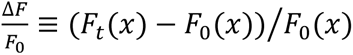. This relative intensity in space and time was averaged over all fish to create a single intensity histogram.

### Principal component analysis (PCA) of speed over time

Tissue speed within 300 µm of the wound centroid was averaged in each frame, and for each larva a track consisting of speed in the first 30 frames (15 minutes) was used for dimensionality reduction. The average speed for all 87 larvae over time was computed and subtracted from each track, and then PCA was performed on the 87 tracks in the 30-dimensional space.

### Non-rigid deformation of LifeAct distributions

Maximum-intensity z-projections of LifeAct in wounded tailfins were registered to remove rigid movement of the entire tissue as described above. A non-rigid warping was applied to further align individual cells, which moved at slightly different speeds in different directions. More explicitly, the goal was to identify warped coordinates (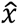, ŷ), so that the fluorescence image *F_t-_*_1_ (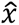(*x*, *y*), ŷ (*x*, *y*) was aligned to the previous frame, *F_t_* (*x*, *y*). To do this, the displacement field *D_t_* (*x*, *y*) = (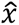, ŷ) was computed, and the frame *t*+1 was warped using those coordinates, so that *F_t-_*_1_ (*D_t_* (*x*, *y*))∼*F_t_* (*x*, *y*), where ∼ indicates similarity in intensity on a pixel-by-pixel basis. The displacement field was computed with the Diffeomorphic Demons algorithm (`imregdemons` in MATLAB with default settings) (Vercauteren et al., 2009). Displacement fields were iteratively composed to register the intensity in each frame to the first frame.

Once movies had been warped to align with the first frame, each frame was smoothed with a Gaussian filter and the relative change in fluorescence intensity at each pixel was computed according to Δ*F/F*_0_ = (*F_t_* (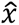, ŷ) – *F*_0_(*x*, *y*)) *F*_0_(*x*, *y*). The average value of Δ*F/F*_0_ was calculated for each frame, excluding the region approximately one cell diameter away from the wound edge. Then these traces of Δ*F/F*_0_ were averaged across all larvae.

Upon inspection, some movies used for motion tracking analysis were not suitable for this analysis of change in LifeAct intensity, due to flickering of illumination light, which led to large frame-to-frame fluctuations in image brightness. Based solely on changes in the background intensity, the following criteria were used to exclude movies used in Fig. 2 A from analysis for Fig. 2 E:

1. The slope of a least-squares fit of average background intensity over time was greater than 0.005×*m* intensity units per frame, where *m* is the median background intensity over the entire movie (typically around 500 intensity units).
2. The average background intensity in the first frame differed by more than 0.5×*m* from *m*, the median background intensity. (Because everything was normalized to the first frame, substantial deviations in the first frame affected the entire trajectory of Δ*F/F*_0_).

### Cell volume measurement

Z-stacks were acquired every 45 seconds at 60x magnification. Subimages of individual cell clusters were manually cropped and deconvolved using the Richardson-Lucy algorithm with 20 iterations in DeconvolutionLab2, a plugin for ImageJ (Sage et al., 2017; Schindelin et al., 2012). A maximum-intensity z-projection of a cell cluster was thresholded to obtain an x-y mask of the cell cluster. To obtain the height of the cell cluster at every other pixel in the x-y mask, the 3D image stack of a cell cluster was smoothed with a 3D gaussian filter and then edges were enhanced with a 3D Sobel filter. Then for a given pixel in the mask, the height was computed by first identifying two peaks in the linescan of fluorescence intensity along the z direction, and then computing the distance between the two peaks. For sub-pixel accuracy in z, the linescans were fitted to gaussians in the vicinity of the peaks. To save on computation time, height was computed at every other pixel. Cell cluster height was spatially smoothed with a 2D median filter and then interpolated to generate a “height map,” the height of the cell cluster as a function of every pixel in the mask of the cluster. The volume of the cell cluster was obtained by numerically integrating this height map using the function `integral2` in MATLAB.

The volume of each cell cluster over time was manually inspected for large discontinuities, and cell clusters for which the height maps had been obviously miscalculated—apparent by large frame-to-frame variations in cell volume over time, as well as visually apparent discontinuities in the height of the cell—were not included for further analysis.

### Code and data availability

Data and code used to generate the figures in this manuscript is available at https://gitlab.com/theriot_lab/fish-wound-healing-nacl

**Figure S1.**
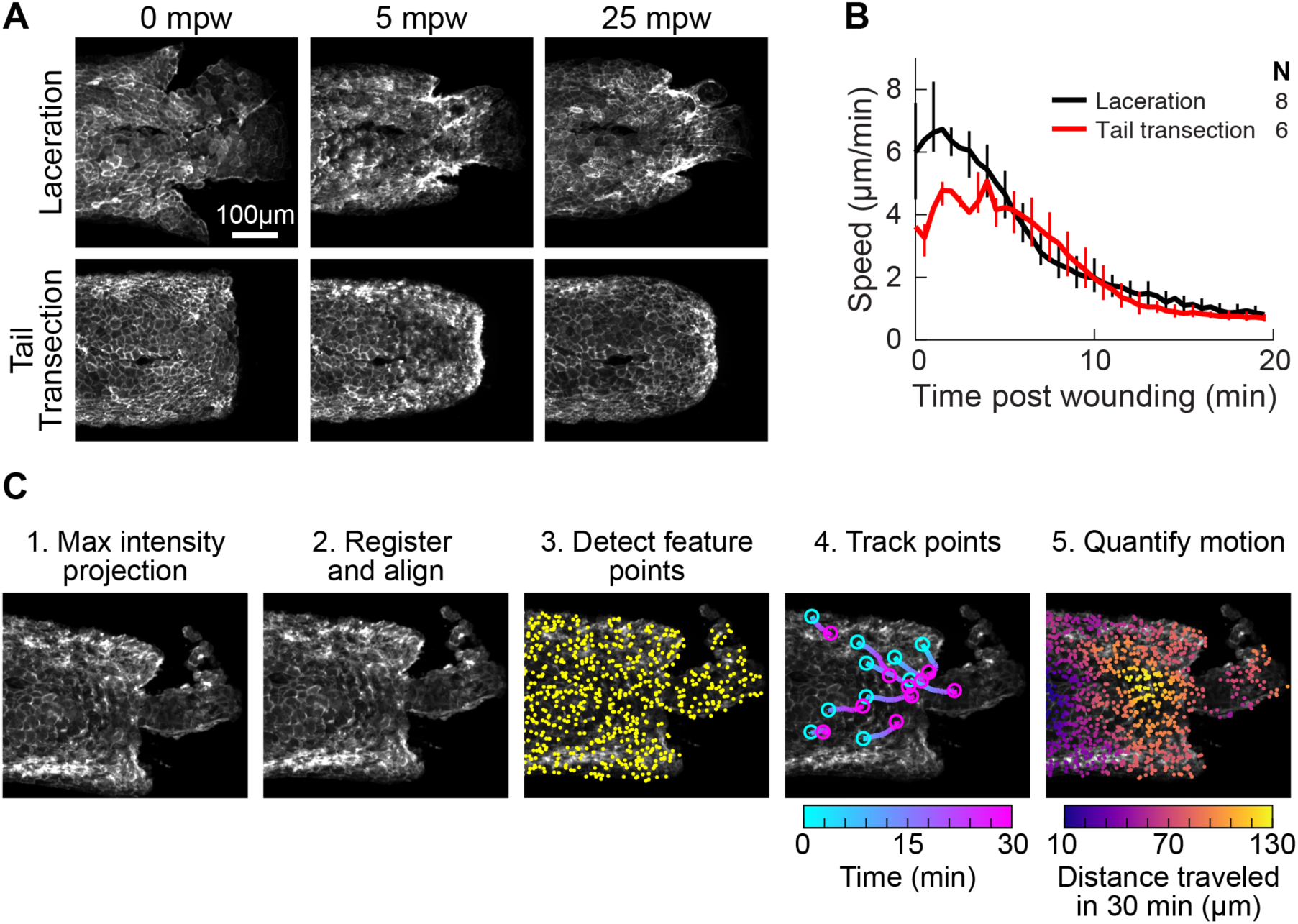
(A) Representative images from timelapse imaging of cellular response to laceration or tail transection. 3 dpf larvae expressing LifeAct-EGFP in basal cells (*TgBAC(ΔNp63:Gal4); Tg(UAS:LifeAct-EGFP); Tg(hsp70:myl9-mApple))* were anesthetized with Tricaine and wounded by laceration or tail transection (see *Methods*) and imaged with spinning disk confocal microscopy. Shown is Z-projections of the LifeAct signal from two representative larvae. (B) Average speed in tissue < 300 µm from the wound for laceration or tail transection. Error bars are bootstrapped 95% confidence intervals. (C) Overview of procedure for tissue motion analysis. 1. Z-projection of LifeAct and myosin light-chain signal. 2. Images of LifeAct are registered using the myosin light chain signal to remove whole-larva drift not due to cell migration. 3. Thousands of feature points are detected throughout the tissue to use as fiducials for motion tracking from frame to frame (for clarity only 15% of the feature points detected in this frame are shown). 4. Feature points are tracked from frame to frame, reporting motion at different locations in the tissue over time. For clarity 10 randomly selected points are shown. 5. Motion is quantified at different positions in the tissue by averaging the movement of feature points across the tissue. Feature points are binned by their distance from the wound, projected along a line extending from the wound centroid anteriorly through the tail.

**Figure S2.**
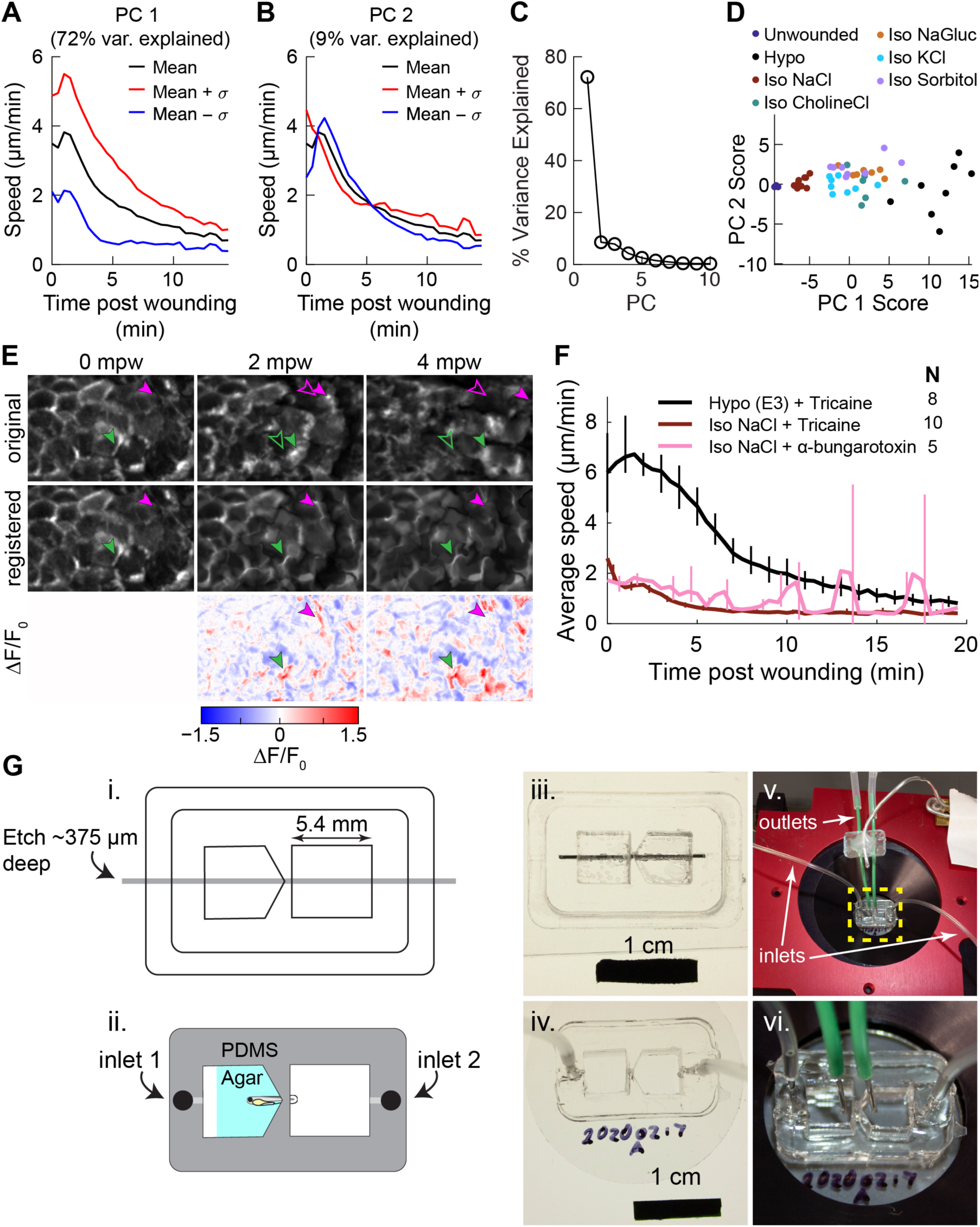
(A-B) Variation along 1st (A) and 2nd (B) principal components of speed trajectories. PC1 captures overall amplitude of the migratory response, while PC2 captures the timing of the peak migratory response. The average trajectory across all conditions is shown in black, and the mean trajectory ± 1 standard deviation along the principal component are shown in red and blue, respectively. (C) Percent variance explained by each of the first 10 principal components. (D) Speed trajectories projected onto the space spanned by the first two principal components. Each dot represents the speed trajectory from one larva. (E) Examples of non-rigid deformation approach to measure changes in actin intensity. Shown is a small region of the LifeAct in basal cells over time, either the original maximum intensity projection (original), the image after non-rigid deformation (registered), and the relative change in fluorescence intensity in the registered image (ΔF/F0). Green and magenta arrowhead show particular LifeAct-rich protrusions; filled arrowheads show the original position of the protrusion, while empty arrowheads show the corresponding position of the protrusion in subsequent frames—which differs due to cell and tissue movement. Registration by non-rigid deformation tracks protrusions and warps the image so all changes in intensity of a protrusion occur at the original location of the protrusion in the first frame. Changes in LifeAct intensity in these protrusions are captured in the ΔF/F0 image at their original position. (F) Speed trajectories for larvae anesthetized with Tricaine or alpha-bungarotoxin. Data for larvae treated with Tricaine is the same as shown in Fig. 2 A. Spikes in the alpha-bungarotoxin condition are due to residual larval twitching due to incomplete muscle relaxation. (G) Two-chamber device schematic. (*i*) Line drawing used in laser cutting pieces of acrylic to make the mold for the two-chamber device. Gray bar indicates the region that was etched rather than cut, to a depth of approximately 375 µm. (*ii*) Schematic of the final device made out of PDMS, with holes punched for fluid inlet. Larvae are immobilized in the device as shown with agar around the anterior part of the fish. (*iii*). Picture of the assembled acrylic mold, with metal bar to create a gap for holding the larvae and for fluid inlets. (*iv*) Photo of the assembled PDMS device cast from the mold, with inlet tubes added. (*v*) Photo of the device in a microscope stage insert with inlet and outlet tubes in place. (*vi*) zoom-in of the device showing the positioning of the inlet and outlet tubes for both chambers to allow independent media exchange in each chamber.

## Notes

https://gitlab.com/theriot_lab/fish-wound-healing-nacl

### Competing Interest Statement

The authors have declared no competing interest.

